# Bootstrap Evaluation of Association Matrices (BEAM) for Integrating Multiple Omics Profiles with Multiple Outcomes

**DOI:** 10.1101/2024.07.31.605805

**Authors:** Anna Eames Seffernick, Xueyuan Cao, Cheng Cheng, Wenjian Yang, Robert J. Autry, Jun J. Yang, Ching-Hon Pui, David T. Teachey, Jatinder K. Lamba, Charles G. Mullighan, Stanley B. Pounds

## Abstract

**Motivation:** Large datasets containing multiple clinical and omics measurements for each subject motivate the development of new statistical methods to integrate these data to advance scientific discovery.

**Model:** We propose bootstrap evaluation of association matrices (BEAM), which integrates multiple omics profiles with multiple clinical endpoints. BEAM associates a set omic features with clinical endpoints via regression models and then uses bootstrap resampling to determine statistical significance of the set. Unlike existing methods, BEAM uniquely accommodates an arbitrary number of omic profiles and endpoints.

**Results:** In simulations, BEAM performed similarly to the theoretically best simple test and outperformed other integrated analysis methods. In an example pediatric leukemia application, BEAM identified several genes with biological relevance established by a CRISPR assay that had been missed by univariate screens and other integrated analysis methods. Thus, BEAM is a powerful, flexible, and robust tool to identify genes for further laboratory and/or clinical research evaluation.

**Availability:** Source code, documentation, and a vignette for BEAM are available on GitHub at: https://github.com/annaSeffernick/BEAMR. The R package is available from CRAN at: https://cran.r-project.org/package=BEAMR.

**Supplementary Information:** Supplementary data are available at the journal’s website.

## Introduction

As omics technologies continue to evolve, increasingly large amounts of data are available for large cohorts of patients. We often have data from multiple omics platforms (e.g., mRNA expression, DNA methylation, proteomics, metabolomics, etc.) as well as clinical data on multiple outcomes (e.g., minimal residual disease [MRD], overall survival [OS], relapse-free survival [RFS], etc.). For example, The Cancer Genome Atlas (TCGA) program has publicly available genomic, epigenomic, transcriptomic, proteomic, and outcome data for 33 cancer types (https://www.cancer.gov/tcga). Similarly, the TARGET (https://cog.cancer.gov/programs/target) and St. Jude Cloud (https://www.stjude.cloud/) [1] databases offer a variety of omics data for pediatric cancers. Much of these data are now available in the Genomic Data Commons (https://gdc.cancer.gov/). These resources present an exciting opportunity to deepen our understanding of the complex biology of genes and their roles in disease. The challenge is how to effectively integrate the multiple forms of omics data to gain clinically valuable insights.

Many multi-omics data integration methods have been developed for dimension reduction and visualization, such as JIVE [2, 3] BIDIFAC [4], iPCA [5], and sparse CCA [6, 7]. Similar methods have been specifically developed for multi-omics single-cell data integration, including MOFA [8], MOFA+ [9], and UMINT [10]. Integrative clustering methods have been developed as well, like intNMF [11], nNMF [12], iCluster [13], iClusterPlus [14], and iClusterBayes [15]. While these methods are useful for exploratory analysis and clustering, they do not directly incorporate outcome data. Some recent methods have been developed to integrate multiple forms of omics data with a single outcome. These methods mainly focus on matrix decomposition and factorization, such as JIVE-predict, where matrix factorization “scores” are included as predictors in models [16] and sJIVE which simultaneously identifies joint and individual components and predicts a continuous outcome [17]. iPCA has also been extended to predict a single clinical outcome, using top PCs as predictors in a random forest model [5]. A Bayesian method, iBAG, uses the underlying biological relationships among molecular features from different platforms to identify genes related to a clinical outcome [18].

Other multi-omics predictive models include those in the *mixOmics* R package, which can integrate multiple omics profiles with a categorical outcome through a variety of dimension reduction techniques and unsupervised or supervised analyses [19]. One such method is DIABLO, which extends sparse generalized canonical correlation analysis to classification problems [20]. LASSO-based predictive models have also been developed, such as the two novel multi-omics variable selection methods to predict cancer prognosis using Cox models [21]. However, these methods have not yet been extended to evaluate multiple clinical outcomes simultaneously.

Many studies still use Venn diagram overlaps to identify genes associated with multiple outcomes at multiple molecular levels (genomic, epigenomic, transcriptomic, proteomic, metabolomic). However, this approach is underpowered [22]. Some studies find significant genes for one platform to generate a gene list to be tested by gene set enrichment analysis (GSEA) for another platform [23]. Still, this approach doesn’t identify individual genes associated with multiple outcomes.

To integrate one form of omic data with multiple clinical outcomes, we have previously developed projection onto the most interesting statistical evidence (PROMISE) [22]. This permutation-based method was shown to have excellent statistical properties and practical value. With very limited cohort sizes, we used PROMISE to successfully identify and validate 60 expression probesets, corresponding to 53 prognostic genes, for childhood acute myeloid leukemia (AML) [24].

We also extended PROMISE to two omics with CC-PROMISE (canonical correlation PROMISE) [25]. We used CC-PROMISE to integrate two forms of omics data to discover that demethylation and overexpression of the methylation writer gene *DNMT3B* are associated with greater total genome-wide methylation and worse prognosis in pediatric AML [26]. This seminal discovery provided the scientific rationale for the ongoing multi-center AML16 clinical trial (clinicaltrials.gov/NCT03164057).

However, PROMISE and CC-PROMISE are limited to evaluating at most two forms of omics data simultaneously and in their ability to adjust for other factors. These methods account for covariates by stratification of the test statistic and stratifying permutation. This can be difficult, especially as the number of covariates grows. When there are too many factors to adjust for, the size of each stratum becomes prohibitively small. Additionally, PROMISE and CC-PROMISE rely on defining the directions of association that are detrimental or beneficial, which is not always straightforward in practice.

Here, we propose the bootstrap evaluation of association matrices (BEAM), a novel multi-omics, multi-outcome, integrative analysis method. BEAM relies on bootstrapping rather than permutation, and thus has some unique capabilities. It allows the evaluation of any number of omics profiles with multiple outcomes. We can evaluate adjusted and unadjusted analyses simultaneously and provide a consensus ranking. Compared to permutation tests, the bootstrap procedure allows for more naturally adjusted analyses.

## Methods

### Notation

For each of *n* = 1, … , *N* subjects, suppose we have collected *C* clinical outcomes (e.g., minimal residual disease [MRD], event-free survival [EFS], and overall survival [OS]) and *k* = 1, … , *K* types of omics data (e.g., methylation, expression, and genotype data). Suppose there are *F*_*k*_ features (e.g., CpG sites, expression probesets, and single nucleotide polymorphisms [SNPs]) for each omic data set *k*. We define sets of these omics features by using their genomic position to map features to gene locations, and call these “gene-feature” sets. Let *s* = 1, … , *S* index the gene-feature sets for which the omics data are available and let *P*_*s*_ index the number of omics features for set *s*. Note that sets can be defined in other ways, such as features belonging to genes in a pathway or located in a particular chromosome arm.

### BEAM

While BEAM can integrate an arbitrary number of omics datasets and clinical outcomes, we will focus on an illustrative example with expression, methylation, and genotype data as omics features, and MRD, EFS, and OS as clinical outcomes (Figure 1). To conduct a BEAM analysis, we first consider the data layout and define the gene-feature sets. For example, in Figure 1, we start with an *N* × *C* matrix of clinical outcomes. Here, *N* = 8 subjects and *C* = 3 for the example outcomes MRD, EFS, and OS. We also have *K* omics datasets each *N* × *F*_*k*_. In this example illustration, *K* = 3 corresponding to genotype data with *F*_1_ = 3 SNPs denoted *G*_1_, *G*_2_, *G*_3_; methylation data with *F*_2_ = 3 CpG sites denoted *M*_1_, *M*_2_, *M*_3_; and transcription data with *F*_3_ = 3 transcripts denoted *T*_1_, *T*_2_, *T*_3_. We define the gene-feature sets by mapping these omics features to two genes based on genomic position. We define the Gene 1 Omics matrix (Set 1), with *N* = 8 rows and *P*_1_ = 4 columns (Figure 1). We also define the Gene 2 Omics matrix (Set 2) with *N* = 8 rows and *P*_2_ = 6 columns. Notice that each set can contain multiple genomic features of the same type and that a single genomic feature (e.g., *G*_2_) can be mapped to multiple sets. In practice, bioinformatic databases such as Ensembl or KEGG can be used to define gene-feature sets based on genomic location or known molecular interactions.

**Figure 1:**
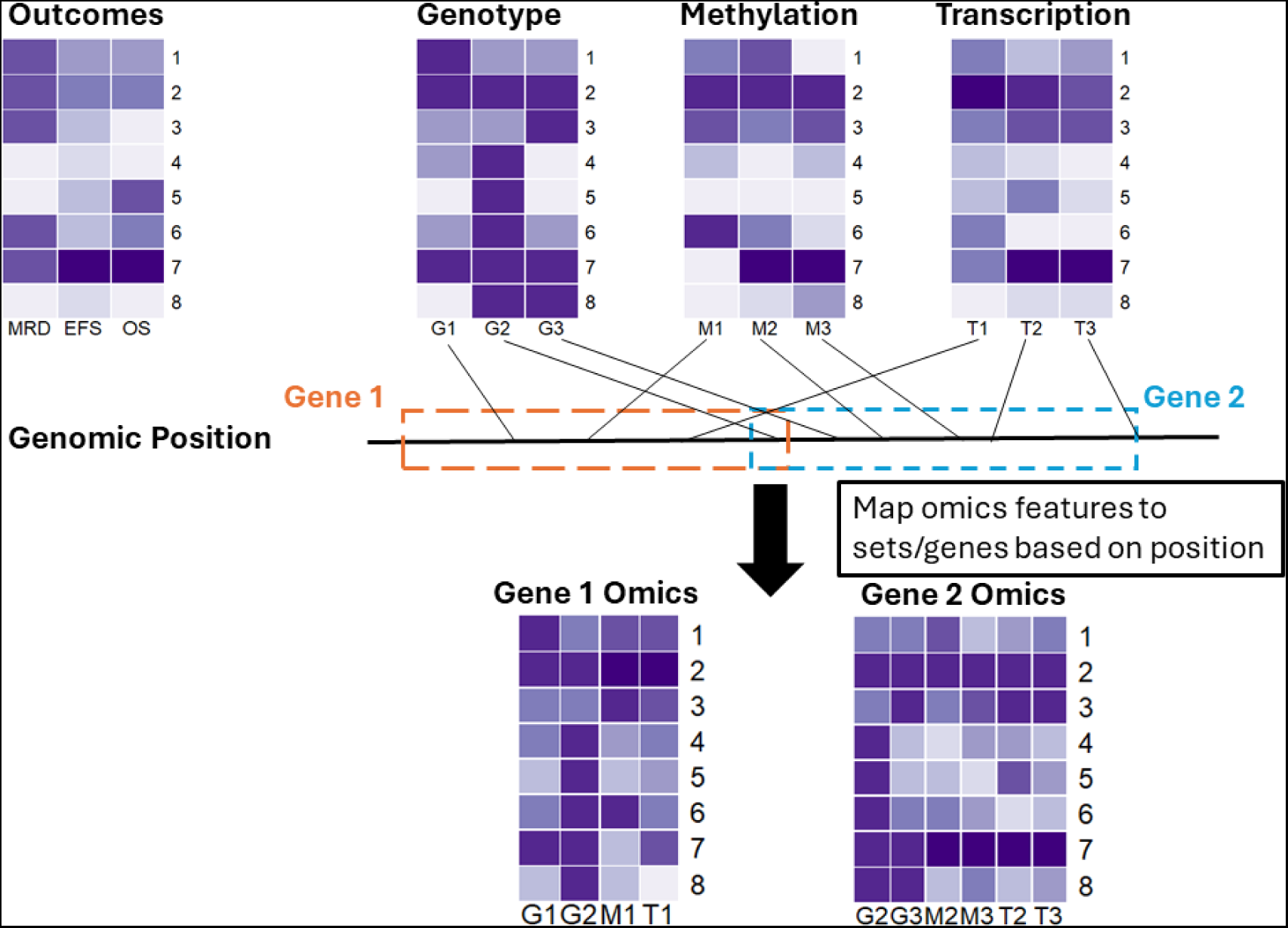
BEAM data layout. Align outcome and omic data matrices. Define gene-feature sets of omic variables by mapping the omics features to genes using genomic position.

Once we have the gene-feature sets defined, we can begin the statistical analysis procedure of BEAM. For a single gene-feature set, we use the outcome matrix and the omics matrix for that set to calculate the association estimate matrix (AEM). For example, in Figure 2, we use the Gene 1 omics matrix (Set 1), which results in an AEM that is *C* × *P*_1_ and shown as the red-blue heat map. Each entry in this AEM is the association found from a regression model fit for each outcome, and each omics feature within the set. For example, in the AEM, the association of a censored event-time outcome with an omic variable can be represented by the regression coefficient from a Cox model using the omic variable as a predictor of the event-time variable (possibly adjusted for covariates). Similarly, logistic and linear regression can be used to obtain coefficients to represent the association of an omic variable with binary and quantitative outcome variables in the AEM, respectively. Next, this AEM is projected into multi-dimensional association estimate space, shown as the pink point in the grey plot. The green point corresponds to the null, that is the point where all the univariate associations are zero (Figure 2a). We use the distance from the observed point to the null (typically the point at which all regression coefficients equal zero) to determine whether the omic features of this set are significantly associated with the clinical outcomes.

**Figure 2:**
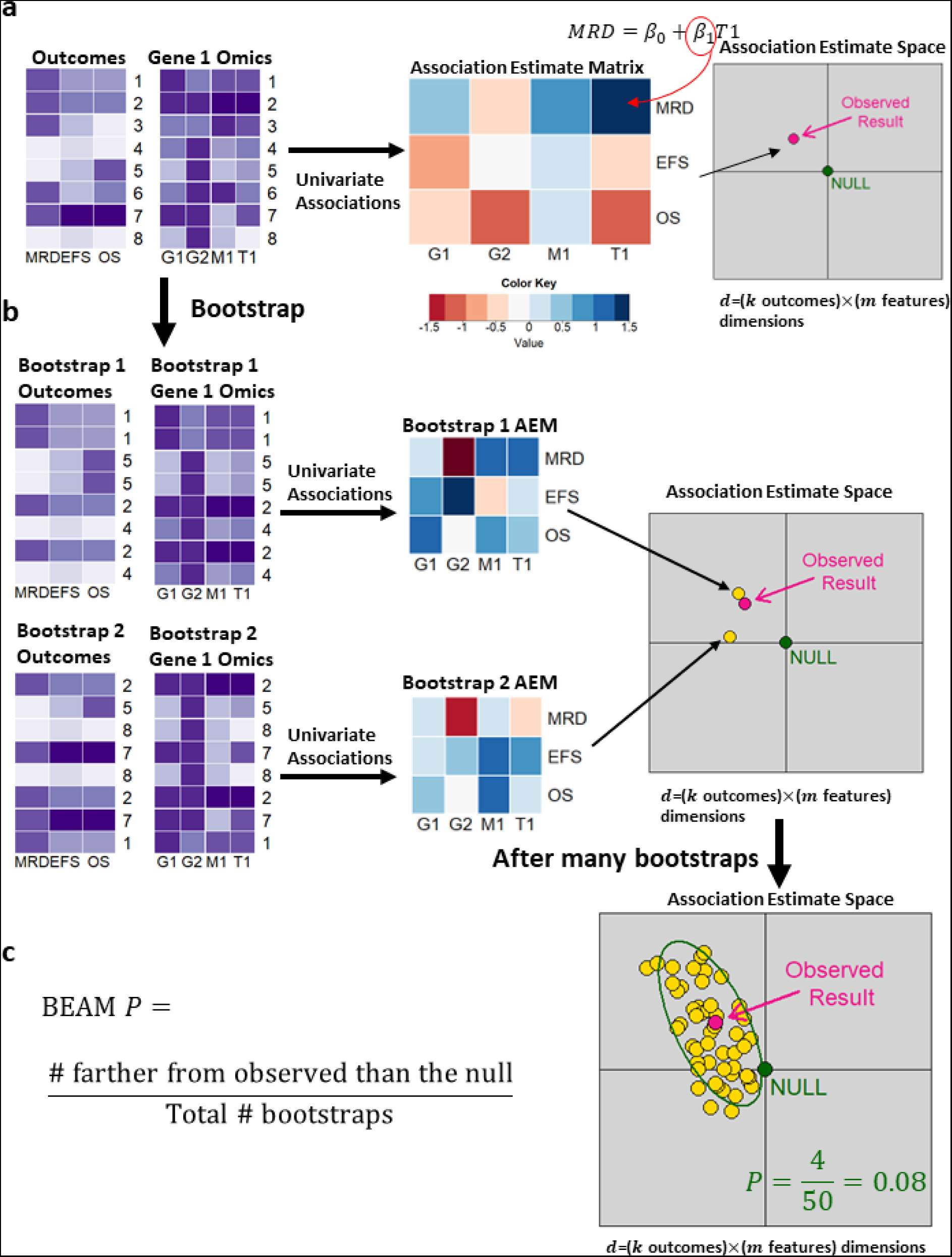
(a) For a gene-feature set, build association estimate matrix (AEM) of regression coefficients from single-feature analyses. Project this observed AEM into multivariate association estimate space (pink) and compare its distance from the green null point of no associations. (b) Bootstrap the cases, maintaining the connection of outcomes and omics features. For each bootstrap resample, construct the AEM and project into multivariate space (yellow points). (c) After many bootstrap resamples, we have a cloud of yellow bootstrap points around the pink observed point. Compute the distance from the observed point of each bootstrap point and the null. Calculate the BEAM P-value.

We use bootstrapping to determine whether the observed point differs significantly from the null. We resample the subject IDs with replacements to form new outcomes and Set 1 omics datasets. Note that we maintain the connection between the omics and the outcome matrices by resampling subjects. For each new bootstrap dataset, we calculate the AEM and again project this as a point in the association estimate space, shown as a yellow point in Figure 2b. We then repeat the bootstrap resampling procedure, resulting in additional points shown in yellow in the association estimate space.

After performing many bootstrap replicates, we have a cloud of bootstrap points (shown in yellow) around the pink observed point (Figure 2c). This cloud of bootstrap points is represented as a *B* × *P*_1_ matrix, as if the bootstraps are observations and the association estimates are variables. We then compute scaled principal components for this matrix, using the observed result vector as the center. In PC space, we compute the Euclidean distance of the null to the observed point and from each bootstrap to the observed. This is equivalent to Mahalanobis distance [27]. The set-level BEAM *P*-value is defined as

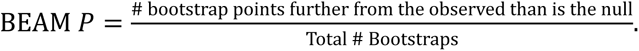

This formula for the p-value is derived by inverting the test technique for confidence interval calculations [28] in the context of empirical bootstrap confidence interval calculations [29]. In other words, we invert the empirical bootstrap confidence interval to obtain a bootstrap *P*-value. The calculation of this *P*-value is illustrated Figure 2c. The green ellipse marks the boundary of the distance from the null point to the observed result. Notice that four bootstrap points fall outside of this ellipse, indicating that it is further from the observed than is the null. Since there are 50 bootstrap points in this example, the BEAM *P*-value is 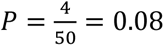. When the observed is far from the null, very few bootstrap points will fall outside of the ellipse, leading to a small *P*-value. When the observed is close to the null, nearly all of the bootstrap points will fall outside of the ellipse, leading to a large *P*-value which indicates a lack of significance (Supplementary Figure 1).

The BEAM procedure is applied to all gene-feature sets, so that the BEAM *P*-value is calculated for all sets. We then use the Pounds-Cheng *q*-value method to account for multiple comparisons [30]. Furthermore, we calculate a distance ratio statistic to evaluate ranking in case of tied *q*- or *P*-values.

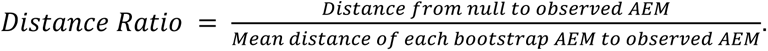

Any number of integrated analyses or simple analyses can be conducted using BEAM. For a set, the AEM can be formed using only features from a particular omics platform, or only using associations with one outcome (Supplementary Figure 2). The AEM could also be formed at the feature level instead. Additionally, a PROMISE-type analysis could be performed if we specify a projection vector of the most interesting associations (see [22]). Then the PROMISE statistic is calculated from the dot product of the z-scaled feature-level AEM and the projection vector (not yet implemented in software).

## Simulations

### Design

We evaluated the performance of BEAM through simulation studies. All simulation studies were conducted in R v. 4.2.0 on the St. Jude Children’s Research Hospital’s high-performance computing facility. Code to implement BEAM is available as an R Package at https://cran.r-project.org/package=BEAMR. Example simulation study code can also be found on GitHub at https://github.com/annaSeffernick/BEAM_Paper.

We used a latent variable approach to generate a variety of null and non-null simulation settings. In each setting, we generated data for one binary outcome, one continuous decimal outcome, and one censored event-time outcome, 10 SNPs, five methylation markers, and two expression transcripts. In the null settings, there were no associations between omics features and outcomes. We also looked at five alternative association structures: (i) 1 SNP associated with all outcomes, (ii) 1 methylation marker associated with all outcomes, (iii) 1 expression probe set associated with all outcomes, (iv) 1 SNP, 1 methylation marker, and 1 expression probe set associated with all outcomes, and (v) all features with all outcomes. Each alternative association structure was simulated with a moderate or a strong effect size. Additionally, we varied the sample size for each setting (*n* = 50, 100, 500, 1000), for a total of 44 simulation settings (4 null settings, one for each of 4 sample sizes; 40 alternative settings defined by 5 association structures x 2 effect sizes x 4 sample sizes). For each setting, we used *B* = 1000 bootstrap replicates and *r* = 1000 simulation replicates. For full details on the simulation study structure, see Supplementary Materials.

BEAM is a very flexible method, and in this simulation study, we fit 33 variations of BEAM for each simulation setting:

- 1 BEAM overall analysis, integrating all omics features with all outcomes.
- 9 BEAM single omic-single outcome analyses, associating all features of an omic type with an outcome.
- 10 BEAM SNP analyses, associating each SNP with all outcomes.
- 5 BEAM methylation analyses, associating each CpG site with all outcomes.
- 2 BEAM expression analyses, associating each expression probe with all outcomes.
- 3 BEAM omic-single outcome analyses, associating all omics features with an outcome.
- 3 BEAM 2 omic analyses, associating all features from 2 omics types with all outcomes.

If there is no further specification, “BEAM” refers to the integrated analysis of all molecular features with all clinical outcomes available for a particular set.

In these BEAM analyses, we fit logistic regression for the binary outcome, linear regression for the continuous outcome, and Cox models for the survival outcome. We compared BEAM to these simple tests of each omic with each outcome. As there were three outcome variables and 17 omic variables, we evaluated a total of 3x17 = 51 simple association tests in our simulations. Additionally, we compared the performance of BEAM to existing integrative methods, PROMISE [22] and CC-PROMISE [25] described in the introduction. We used the R packages *PROMISE* and *CCPROMISE*, available on Bioconductor. PROMISE results are comparable to the BEAM analyses associating a genomic feature with all outcomes, and the CC-PROMISE analyses are comparable to the BEAM 2 omic analyses. Finally, we compared BEAM to two single omics integrative gene set methods: sequence kernel association test (SKAT) [31] and the global test [32]. SKAT evaluates the association of sets of SNPs with a single outcome through kernel machine regression [31, 33–36] and was implemented using the *SKAT* R package. SKAT has also been extended to survival outcomes [37], which is implemented in the *seqMeta* package available on GitHub (https://github.com/hanchenphd/seqMeta). The global test was designed to test the association of expression of groups of genes with a binary, continuous, or survival clinical outcome [32, 38]. As the global test is based on a random effects model, it can be applied to methylation and genotype data as well. We used the R package *globaltest* in our simulations. These tests are comparable to the BEAM single omic-single outcome analyses, which integrate possibly multiple omics features of the same type with a single outcome.

### Results

Simulation results can be found in the Supplementary Materials. Table S1 provides details for all simulation settings including the sample size, effect size, and the associated coefficient matrix *M*. Table S2 provides the mean *P*-value, Pr(*P* < α) for α = 0.01, 0.05, and purity for each analysis performed on each simulation setting. Purity is the proportion of true non-zero associations for a gene-feature set and collection of outcomes. For the null settings, the purity is zero, and for the settings where all features are associated with all outcomes, the purity is one.

In the null datasets, where none of the omic features are associated with the clinical outcomes (Settings 1-4, Tables S1-S2), BEAM maintains the nominal Type I error rate. In the alternative settings (Settings 5-44, Tables S1-S2), BEAM generally performs better in terms of greater statistical power as the sample size increases and the number of features associated with the outcomes increases. In Table 1, the top methods with greatest power and smallest mean *P*-value are reported for each simulation setting with sample size n=100 and moderate effect size (d=0.5). At least one BEAM analysis variation is in the top three methods for each setting, and similar results are observed for the other settings (Table S3). We call the univariate test with the greatest power the “best simple test.” For example, in Table 1, the best simple test for setting 6, which has one truly associated SNP, is the simple test of this SNP (labeled gtyp1) with the decimal (continuous) outcome. However, the best simple test would not be known in practice, as we don’t know which genomic features are truly associated with the outcomes of interest in real data. Fortunately, BEAM analyses often have power similar to that of the best simple test.

**Table 1:** Top 3 methods for each alternative setting with sample size n=100 and effect size d=0.5.

| Setting | Associated Omic | Analysis Method (omic, outcome) | Power (0.01) |
| --- | --- | --- | --- |
| 6 | SNP1 | Simple (SNP1, decimal) | 0.224 |
| 6 | SNP1 | BEAM (SNP1, all) | 0.211 |
| 6 | SNP1 | PROMISE (SNP1, all) | 0.091 |
| 14 | Meth1 | Simple (Meth1, decimal) | 0.38 |
| 14 | Meth1 | BEAM (Meth1, all) | 0.368 |
| 14 | Meth1 | Simple (Meth1, binary) | 0.142 |
| 22 | Expr1 | Simple (Expr1, decimal) | 0.456 |
| 22 | Expr1 | BEAM (Expr1, all) | 0.417 |
| 22 | Expr1 | BEAM (Expr, decimal) | 0.297 |
| 30 | SNP1, Meth1, Expr1 | Simple (Expr1, decimal) | 0.42 |
| 30 | SNP1, Meth1, Expr1 | BEAM (Expr1, all) | 0.394 |
| 30 | SNP1, Meth1, Expr1 | Simple (Meth1, decimal) | 0.38 |
| 38 | All | BEAM (all, decimal) | 0.697 |
| 38 | All | BEAM (Meth & Expr, decimal) | 0.697 |
| 38 | All | BEAM (all, all) | 0.693 |

BEAM is a very flexible method, and in this simulation study, we fit several variations of BEAM for each simulation setting. Table 2 shows the top BEAM methods in terms of greatest power and smallest mean *P*-value for each simulation setting with sample size n=100 and moderate effect size (d=0.5). Consistently, the BEAM variation that tests the true association has the greatest power, as expected. For example, in setting 6 with one SNP (labeled gtyp1) truly associated with all outcomes, the BEAM test of this SNP with all outcomes has the greatest power, followed by BEAM tests that involve all SNP (labeled gtyp) variables. We see similar patterns for the other settings in Table 2 and all settings in Table S4. These results show that care must be taken when selecting the type of BEAM analysis to perform. The overall integration of all omics with all outcomes [BEAM (all, all)] may not have the greatest power in all application scenarios. A summary of all simulation settings can be found in Supplementary Figure 5, which shows that a BEAM analysis is in the top three analyses with greatest power for most settings, and that power improves for BEAM and the other integrated analysis methods as sample size and effect size increase.

**Table 2:** Top 3 BEAM methods for each alternative setting with sample size n=100 and effect size d=0.5.

| Setting | Associated Omic | Analysis Method (omic, outcome) | Power (0.01) |
| --- | --- | --- | --- |
| 6 | SNP1 | BEAM (SNP1, all) | 0.211 |
| 6 | SNP1 | BEAM (SNP, decimal) | 0.029 |
| 6 | SNP1 | BEAM (SNP & Expr, decimal) | 0.027 |
| 14 | Meth1 | BEAM (Meth1, all) | 0.368 |
| 14 | Meth1 | BEAM (Meth, decimal) | 0.08 |
| 14 | Meth1 | BEAM (Meth, all) | 0.054 |
| 22 | Expr1 | BEAM (Expr1, all) | 0.417 |
| 22 | Expr1 | BEAM (Expr, decimal) | 0.297 |
| 22 | Expr1 | BEAM (Expr, all) | 0.237 |
| 30 | SNP1, Meth1, Expr1 | BEAM (Expr1, all) | 0.394 |
| 30 | SNP1, Meth1, Expr1 | BEAM (Meth1, all) | 0.371 |
| 30 | SNP1, Meth1, Expr1 | BEAM (Expr, decimal) | 0.292 |
| 38 | All | BEAM (all, decimal) | 0.697 |
| 38 | All | BEAM (Meth & Expr, decimal) | 0.697 |
| 38 | All | BEAM (all, all) | 0.693 |

## An Application Example: Pediatric B-cell Acute Lymphoblastic Leukemia (B-ALL)

For the application analysis, BEAM analyses were conducted in R-4.3.1 on St. Jude high performance computing cluster. Table and figure creation were performed in R-4.2.0. The code of this application is available on GitHub (https://github.com/annaSeffernick/BEAM_Paper).

### Data and BEAM Analyses

We applied BEAM to a multi-omics pediatric B-ALL data set of 170 patients from TOTAL XV (NCT00137111) and TOTAL XVI (NCT00549848) clinical trials who were treated at St. Jude [39]. Most patients had gene expression, measured with Affymetrix HG-U133 arrays; DNA methylation, measured with Illumina 450K array; germline genotypes, measured with Affymetrix Mapping 6.0 or 500KSNP array; and somatic Copy Number Variation (CNV) data derived from the SNP arrays (Supplementary Figure 6). We integrated these four omics profiles with five outcomes: dichotomous MRD at protocol day 19 (middle of remission induction) and day 46 (end of remission induction), continuous LC_50_ of prednisolone (*log*_10_-transformed; dose of prednisolone required to kill 50% of patient leukemic cells *ex vivo*), EFS, and OS. We applied BEAM with Firth-penalized logistic regression [40] for MRD at both time points, linear regression for log(LC_50_), and Firth-penalized Cox regression [41] for EFS and OS, using 1000 bootstrap replicates. Firth-penalization stabilizes regression coefficients for analyses involving small sample sizes or small number of events [40, 41]. Gene-feature sets were defined based on Ensembl ID and genomic position [42]. For SNPs and CpG sites (methylation data), we mapped a feature to a gene-feature set if the feature was within 50kb of the gene’s start and end position. An automated PubMed literature search was performed to annotate the top genes from this analysis. We also explored the ability of BEAM to adjust for additional covariates. We applied BEAM with the same models as described above, except we additionally included leukemia molecular subtype as a categorical variable in each regression model. We again used 1000 bootstrap replicates.

This dataset contains 50,353 gene-feature sets. BEAM identified 157 gene-feature sets with *q* < 0.2 (Table S5), including 26 known leukemia genes identified by an automated PubMed literature search (Table S6). The BEAM analysis found several genes known to be associated with leukemia in the literature, such as *PLAGL2*, *CD27*, and *NOTCH1*; these genes were not identified in the original analysis of this dataset [39]. An adjusted BEAM analysis was also performed, in which each feature-outcome regression model also included leukemia molecular subtype as a covariate. The minimum *q*-value from this analysis was 0.804. However, of the 157 gene-feature sets identified in the unadjusted BEAM analysis, all had this minimum *q*-value and 87 had *P* < 0.05 in the adjusted analysis (Table S7).

One interesting gene identified in the unadjusted BEAM analysis was *CD1C*, a gene that has been implicated in other leukemias [43–45] but was not found in univariate screening or by a customized p-value aggregation method developed for analysis of this dataset in [39]. The p-value aggregation analysis integrated six forms of molecular omics data with the LC_50_outcome. *CD1C* ranked third in this paper’s CRISPR knockout screen (see Supplementary Table 6 in [39]) strongly indicating that it may play a role in glucocorticoid resistance. Chronic B-cell leukemia cells may improve their survival advantage by suppressing the expression of *CD1C* to reduce their interaction with immune cells [43]; also, human T-cells are able to target CD1C+ acute B-cell leukemia cells [44]. Additionally, research suggests *CD1C* is prognostically important in breast cancer [46], cervical cancer [47], and neuroblastoma [48] and also implicated in cancer-immune system interaction [43, 44, 46].

Clinical plots (Figure 3), bootstrap plots (Supplementary Figure 7), and individual association test results suggest that SNPs, expression, and methylation are driving the BEAM significance for *CD1C*. Expression of probeset 205987_at was positively associated and methylation of CpG cg04574507 was negatively associated with log(LC_50_), but these features were not significantly associated with survival or MRD. SNP_A-2076774 was significantly associated with OS and MRD at day 46, while SNP_A-8578231 was significantly associated with EFS. The *CD1C* gene remained significant in the BEAM analysis adjusting for leukemia molecular subtype (*P* = 0.049; Table S7). A table of genotype by subtype for the SNPs that map to *CD1C* can be found in Table S8. Some additional genes present in the CRISPR screens of [39] that were identified by BEAM but not the original integrated analysis are *GYPE*, *CCDC114*, *ARHGAP18*, *MAGI3*, *PARP8*, and *STRADA*.

**Figure 3:**
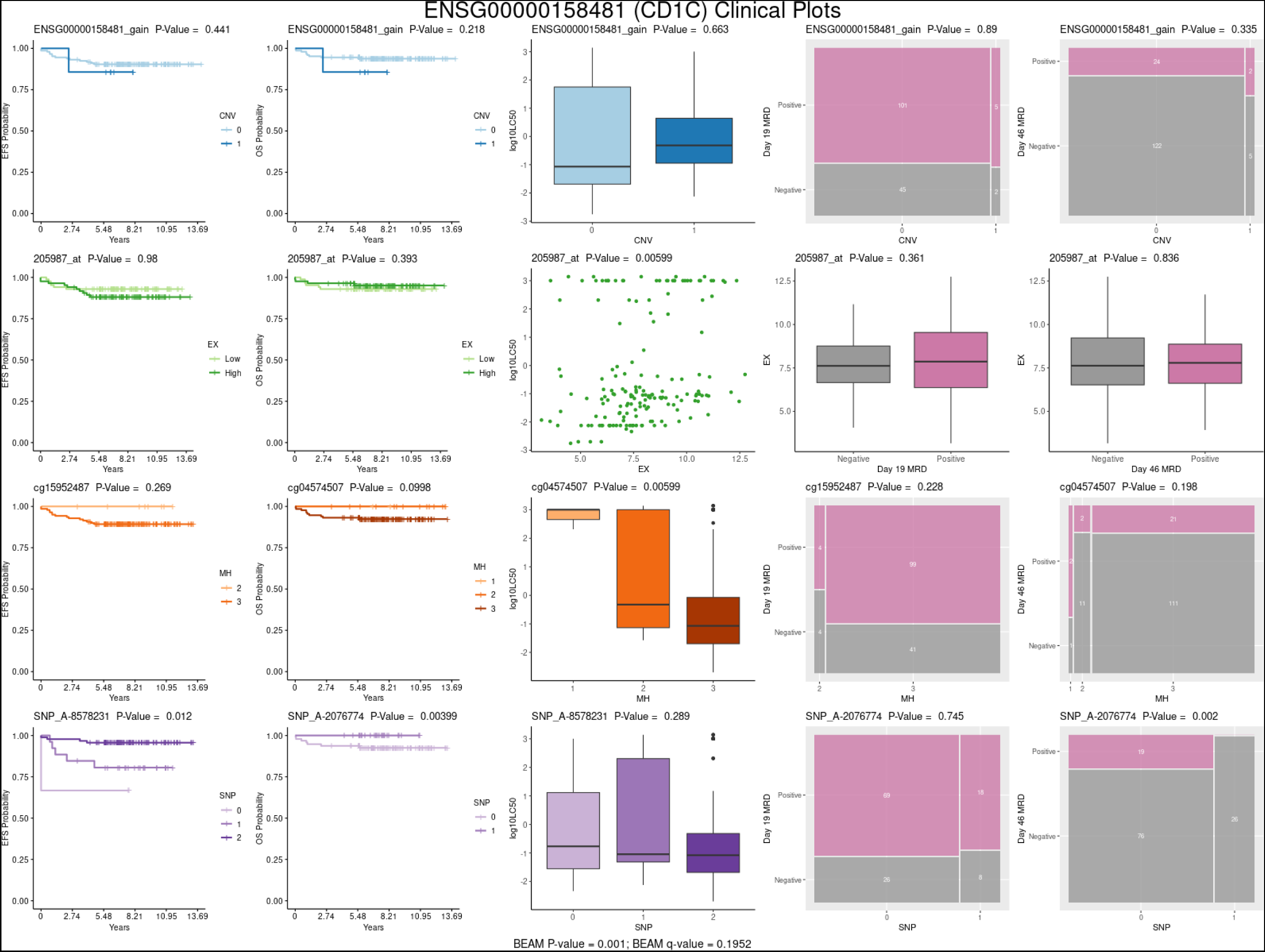
Clinical plots for *CD1C* from BEAM application to TOTAL pediatric B-ALL dataset.

## Discussion

As large datasets containing multiple forms of molecular omics data and multiple clinical outcomes become publicly available, integrated statistical analysis methods are paramount to inform biologically meaningful discoveries. Here, we propose a bootstrap-based integrated analysis method called BEAM that can evaluate the associations of multiple omic variables with multiple clinical outcomes. This method is implemented in an R Package called “BEAMR” available on GitHub (https://github.com/annaSeffernick/BEAMR) and CRAN (https://cran.r-project.org/package=BEAMR). In our simulations and applications, BEAM outperformed other methods in most scenarios. BEAM also maintained type I error rate in null simulation settings and often had the greatest or second-greatest power in alternative settings.

BEAM also performed well when applied to a pediatric B-ALL dataset. This application demonstrated the novelty of BEAM, as it was able to integrate four omics variables with five clinical outcomes, a feat that existing methods could not achieve. BEAM identified both known leukemia-related genes and novel genes, including *CD1C* which had not been previously implicated in pediatric B-ALL and was not found in univariate screens or by another integrated analysis method in the original data analysis. This gene could be an important prognostic biomarker or immunotherapy target [49] in pediatric B-ALL and warrants further studies. Furthermore, CRISPR assays provide experimental evidence that *CD1C* is functionally involved in prednisolone resistance [39].

In addition to integrating an arbitrary number of omics with multiple outcomes, BEAM can also easily incorporate additional covariates. The association estimates in the AEM can be derived from regression coefficients of the omics features in multivariate linear regression models that adjust for confounders or important clinical factors, such as age and sex. Another advantage of BEAM over PROMISE and CC-PROMISE is that BEAM does not require the user to specify a projection vector that defines the direction of associations of interest. This flexibility allows for identifying genes that may be beneficially associated with some outcomes but detrimentally associated with other outcomes. However, if a PROMISE-type analysis is desired, a projection vector can be provided (this capability is not yet implemented in the software).

BEAM is also a very flexible and general method that can be used for various types of integration. After each of the omic/outcome association statistics are calculated, it is straightforward to calculate the integrated BEAM *P*-value for any combination of features and outcomes of interest. Another aspect of flexibility is the type of association statistic that can be input into the BEAM framework. We used regression coefficients, but correlations or even measures of predictive ability could be used instead. This might require reformulating the null hypothesis. Since BEAM was developed based on regression coefficients, the null is defined as a vector of zeros. Other statistics with non-zero nulls could be accommodated, perhaps by applying a transformation first. Incorporating different statistics into the BEAM framework is an intriguing area for future work.

As with other integrated analysis methods, BEAM improves statistical power by combining information across omics datasets. BEAM computes an empirical p-value as the proportion of bootstrap association estimate matrices (AEMs) that are farther from the observed AEM in Mahalanobis distance than the complete null (where no omic variable associates with any outcome variable). One area of future research is to evaluate the use of these components to define weights, allowing certain associations to be prioritized. Additional research directions include improving computational performance to decrease the computation time, incorporating other types of outcomes (e.g., toxicity, adverse event) in addition to efficacy outcomes, and applying BEAM to other high-dimensional data types such as imaging data.

## Competing Interests

There were no direct competing interests related to this work. C-H.P. receives personal fees from Novartis. D.T.T. received research funding from Neoimmune Tech and BEAM Therapeutics (unrelated to the BEAM method described in this manuscript) and serves on advisory boards for BEAM Therapeutics, Sobi, Janssen, Jazz and Servier. D.T.T. has patents or patents pending on CAR-T. C.G.M. serves on the scientific advisory board and receives honoraria for Illumina, and received research funding from Pfizer, equity from Amgen and royalties from Cyrus. J.J.Y. receives research funding from Takeda Pharmaceutical Company and AstraZeneca plc. X.C, J.K.L, and S.B.P have a patent for Pharmacogenomics Score to Make Decisions on Therapy Augmentation in AML pending 18/683,969. S.B.P also receives grants from Gateway for Cancer Research and the American Cancer Society and has patents pending for Leukemia Diagnostic Based on Gene Expression, Methods for Predicting AML Outcome, and AML Risk Stratification Using OS iScore.

## Acknowledgements

The authors gratefully acknowledge the St. Jude high performance computing (HPC) facility staff for their help implementing this method on the HPC, Lakshmi Anuhya Patibandla for annotating our gene list with PubMed citations, and Lei Shi for his help with early implementations of this method.

## Funding

This work was supported by St. Jude Children’s Research Hospital; American Lebanese Syrian Associated Charities (ALSAC); the National Institutes of Health National Cancer Institute [R01CA132946 (JKL, SBP), R01CA270120 (JKL, SBP), T32CA236748 (CGM), CA021765 (Cancer Center Core Grant)]; and the National Institutes of Health Gabriella Miller Kids First Pediatric Research Program [X01HD100702 (CGM, DT)].

## Data Availability

Simulation data are available on GitHub (https://github.com/annaSeffernick/BEAM_Paper). Gene expression and DNA methylation data for the pediatric B-ALL example are available at Gene Expression Omnibus under accession no. GSE66708 (https://www.ncbi.nlm.nih.gov/geo/query/acc.cgi?acc=GSE66708). Genotype data is available upon request at dbGaP (https://www.ncbi.nlm.nih.gov/projects/gap/cgi-bin/study.cgi?study_id=phs000638.v1.p1).

## Supplementary Materials

Supplementary materials are listed below and available in the online version of this article. Supplementary analysis codes are available online at https://github.com/annaSeffernick/BEAM_Paper.

**Supplementary** Figure 1: Beam P-value explanation.

**Supplementary** Figure 2: Different types of BEAM analyses.

**Supplementary** Figure 3: Schematic of simulation design.

**Supplementary** Figure 4: Illustration of latent variable data generation approach.

**Supplementary** Figure 5: Simulation Summary.

**Supplementary** Figure 6: UpSet plot of B-ALL application data.

**Supplementary** Figure 7: Bootstrap plot from B-ALL application.

**Supplementary Table S1:** Simulation Settings.

**Supplementary Table S2:** Simulation Results.

**Supplementary Table S3:** Top 3 methods for each simulation scenario.

**Supplementary Table S4:** Top 3 BEAM variations for each simulation scenario.

**Supplementary Table S5:** BEAM analysis results of pediatric B-ALL application.

**Supplementary Table S6:** Literature annotation results of top BEAM findings in pediatric B-ALL application.

**Supplementary Table S7:** Adjusted BEAM analysis results of B-ALL application.

**Supplementary Table S8:** Cross tabulation of genotype and subtype for SNPs that map to *CD1C*.

## Author Contributions

Conceptualization: S.B.P., X.C., C.C., J.K.L.; methodology: A.E.S., S.B.P., X.C., C.C.; software: A.E.S., S.B.P., X.C.; simulation studies: A.E.S., S.B.P., X.C.; data acquisition: W.Y, R.J.A., J.J.Y, C-H.P., C.G.M.; data analysis: A.E.S., W.Y., S.B.P.; analysis interpretation: A.E.S., S.B.P., J.K.L., D.T.T., C.G.M., writing-review & editing: all authors; writing-original draft: A.E.S., S.B.P.; supervision: S.B.P.; funding acquisition: J.K.L., S.B.P., C.G.M., D.T.T.

## Notes

https://www.ncbi.nlm.nih.gov/geo/query/acc.cgi?acc=GSE66708

https://www.ncbi.nlm.nih.gov/projects/gap/cgi-bin/study.cgi?study_id=phs000638.v1.p1

https://github.com/annaSeffernick/BEAM_Paper

