## Supplementary Materials for "Bootstrap Evaluation of Association Matrices (BEAM) for Integrating Multiple Omics Profiles with Multiple Outcomes"

[1]Department of Biostatistics, St. Jude Children's Research Hospital, Memphis, TN, USA; [2]Department of Health Promotion and Disease Prevention, University of Tennessee Health Science Center, Memphis, TN, USA; [3]Department of Pharmacy & Pharmaceutical Services, St. Jude Children's Research Hospital, Memphis, TN, USA; [4]Hematological Malignancies Program, St. Jude Children's Research Hospital, Memphis, TN, USA; [5]Hopp Children's Cancer Center Heidelberg (KiTZ), Heidelberg, Germany; [6]Division of Pediatric Neurooncology, German Consortium for Translational Cancer Research (DKTK), German Cancer Research Center (DKFZ), Heidelberg, Germany; [7]Department of Oncology, St. Jude Children's Research Hospital, Memphis, TN, USA; [8]Department of Pathology, St. Jude Children's Research Hospital, Memphis, TN, USA; [9]Perelman School of Medicine at the University of Pennsylvania, Philadelphia, PA, USA; [10]Department of Pediatrics and the Center for Childhood Cancer Research, Children's Hospital of Philadelphia, Philadelphia, PA, USA; [11]Division of Oncology and Center for Childhood Cancer Research, Children's Hospital of Philadelphia, Philadelphia, PA, USA; [12]Department of Pharmacotherapy and Translational Research, University of Florida College of Pharmacy, Gainesville, FL, USA

### Supplementary Methods

**Supplementary Figure 1:** The BEAM P-Value is calculated using Mahalanobis distance. The green ellipse corresponds to the distance from the observed result to the null. When the observed is far from the null (panel A), no bootstrap points fall outside of the ellipse, leading to a small p-value. In panel B, the observed is close to the null so most bootstraps fall outside of the ellipse. This leads to a large P-Value.

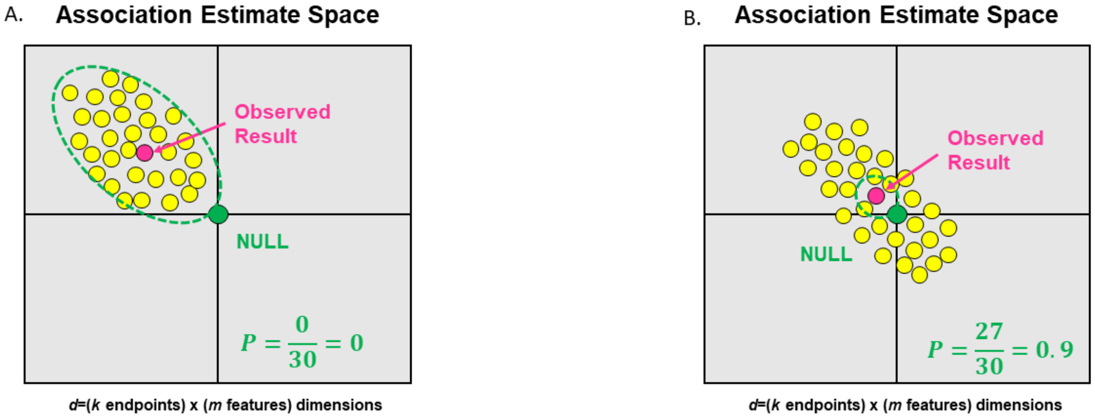


$$\text{BEAM } P = \frac{\text{\# bootstraps farther from the observed than is the null}}{\text{Total \# Bootstraps}}.$$

**Supplementary Figure 2:** BEAM can perform a suite of complimentary integrated tests. BEAM can be used to associate any number of omics with any number of endpoints.

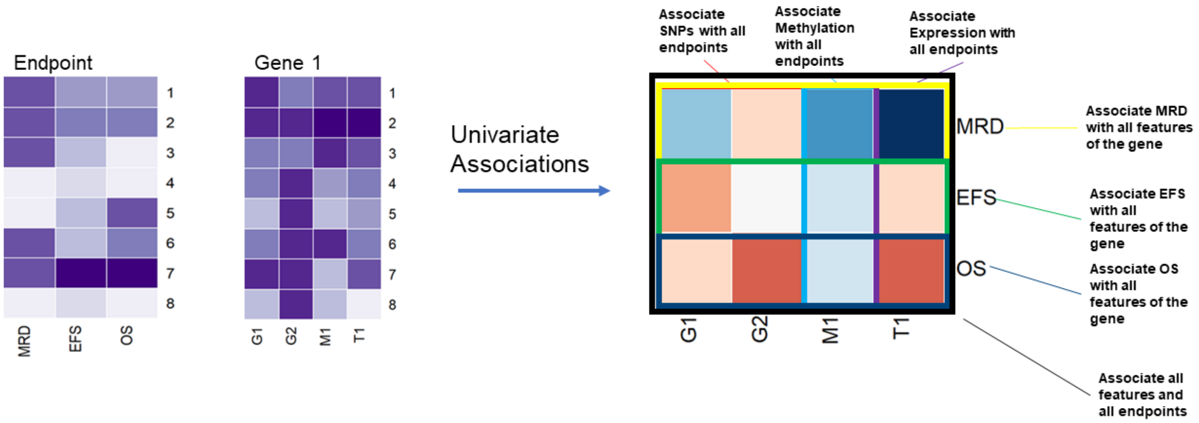

**Supplementary Simulations**

**Supplementary Figure 3: Schematic of Simulation Design.**

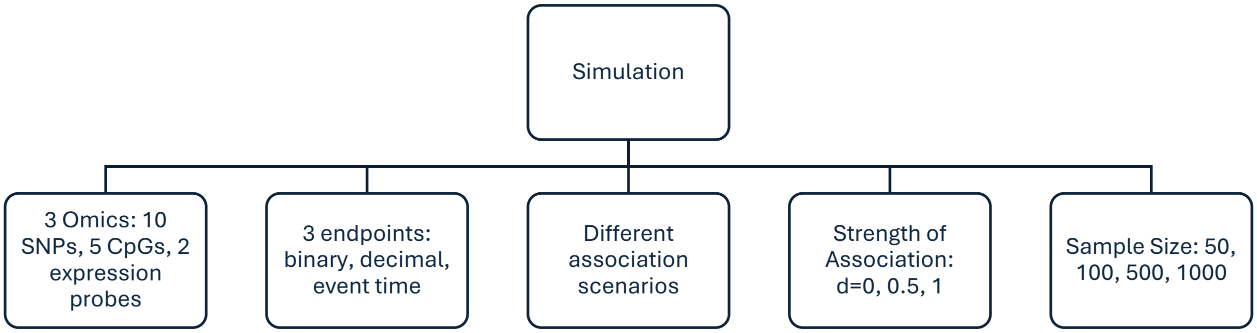


We simulated data with 3 endpoints and 3 omics. We used a latent variable approach to generate a single gene with 10 SNPs, 5 CpG sites and 2 expression probes (Supplementary Figure 4). We considered different scenarios of association: the null setting where none of the omics were associated with endpoints, the case were a single feature of one omic was associated with all endpoints (e.g., SNP1 associated with all endpoints); case where 1 feature of each omic type was associated with all endpoints (e.g., SNP1, Meth1, Expr1); finally, case where all omics features were associated with all endpoints.

Transformation of Latent Scores to Analysis Dataset:

- Binary Endpoints: $y \sim Bernoulli(\Phi(x))$
- Continuous Quantitative Endpoints: $y = x$
- Survival Time Endpoints (like a SJ clinical trial):
    - Censoring time: $t_c \sim Uniform(3,8)$
    - Event time: $t_e \sim Exponential\left(1.05 - \Phi(x)\right)$
    - Observed time: $t_o = min(t_c, t_o)$
    - Observed event status: $\mathrm{e_0} = \mathrm{I}(\mathrm{t_e} < \mathrm{t_c})$
- Expression: $y = x$
- Methylation: $y = \Phi(x)$
- Genotype: $y \sim Binomial(2, \Phi(x))$

We varied the strength of association from no association, weak association, and stronger association, $d$=0, 0.5, 1. We also varied the sample size $n$=50, 100, 500, 1000. This results in 44 total simulations settings. We compared the performance of BEAM with other integrated analysis methods: PROMISE, CC-PROMISE, SKAT, and the global test. In null simulation settings we evaluated type I error and in alternative settings we compared power.

**Supplementary Figure 4: Visualization of latent variable approach to generating simulation data.**

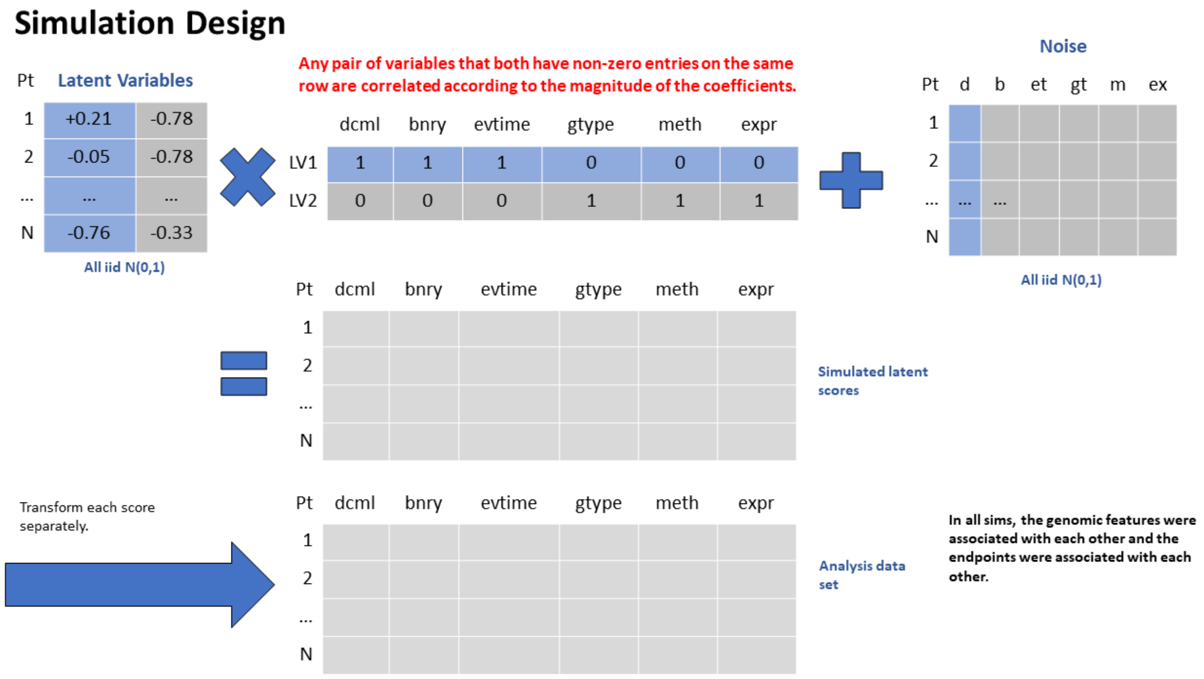

**Supplementary Figure 5: Summary of all simulation settings in terms of power at $\alpha = 0.01$.**

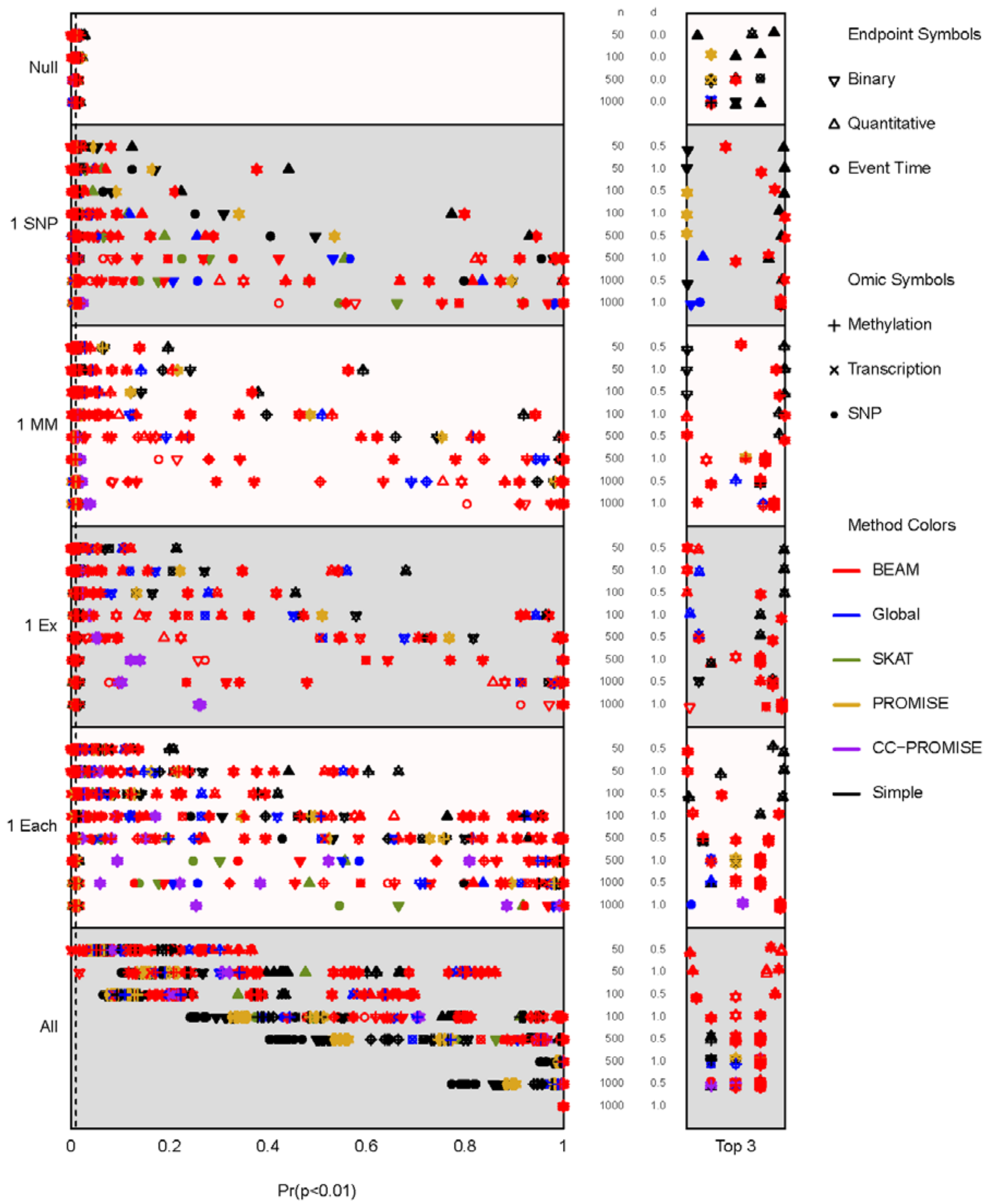

### Supplementary Application

**Supplementary Figure 6:** Upset plot of available omics data from TOTAL XV and TOTAL XVI pediatric B-ALL St. Jude dataset. SNP=genotype data, MH=methylation data, CNV=copy number variation data, EX=gene expression data. 120/170 patients had all four types of omics data.

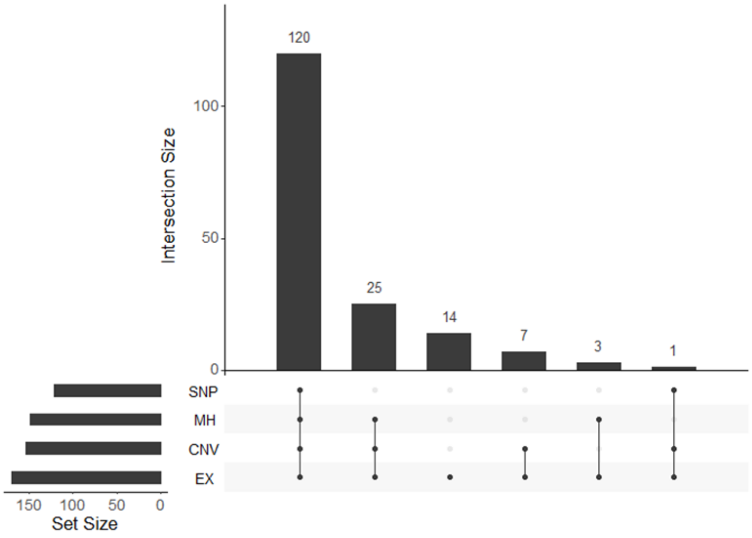


**Supplementary Figure 7:** Example bootstrap plots for *CD1C*, found by BEAM application to pediatric B-ALL dataset. The observed point is in coral, null is in teal, and bootstraps are grey circles. The black crosses correspond to bootstrap replicates that were further from the observed than is the null.

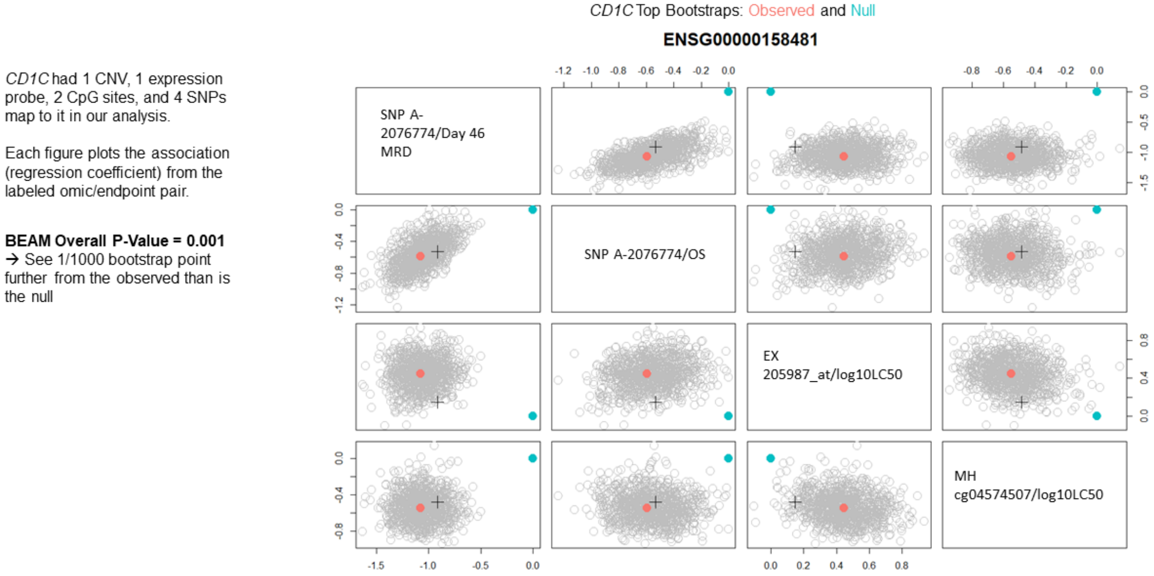
